# SERINC5 potently restricts retrovirus infection *in vivo*

**DOI:** 10.1101/2020.05.19.102798

**Authors:** Uddhav Timilsina, Supawadee Umthong, Brian Lynch, Aimee Stablewski, Spyridon Stavrou

## Abstract

The Serine Incorporator (SERINC) proteins are multipass transmembrane proteins that affect sphingolipid and phosphatidylserine synthesis. Human SERINC5 and SERINC3 were recently shown to possess antiretroviral activity to a number of retroviruses including human immunodeficiency virus (HIV), murine leukemia virus (MLV) and equine infectious anemia virus (EIAV). In the case of MLV, the glycosylated Gag (glyco-Gag) protein was found to counteract SERINC5-mediated restriction in *in vitro* experiments and that the viral envelope determines virion sensitivity or resistance to SERINC5. However, nothing is known about the *in vivo* function of SERINC5. Antiretroviral function of a host factor *in vitro* is not always associated with antiretroviral function *in vivo*. Using SERINC5-/- mice we generated, we show that mouse SERINC5 (mSERINC5) restriction of MLV infection *in vivo* is dependent not only on glyco-Gag, but also on the retroviral envelope. Finally, we also examined the *in vivo* function of the other SERINC gene with known antiretroviral functions, SERINC3. By using SERINC3-/- mice, we found that the murine homologue, mSERINC3, had no antiretroviral role both *in vivo* and *in vitro*. This report provides the first data showing that SERINC5 restricts retrovirus infection *in vivo* and that restriction of retrovirus infectivity *in vivo* is dependent on both the presence of glyco-Gag and the viral envelope.

**IMPORTANCE:** This study examines for the first time the *in vivo* function of the <u>Ser</u>ine <u>Inc</u>orporator (SERINC) proteins during retrovirus infection. SERINC3/5 restrict a number of retroviruses including human immunodeficiency virus 1 (HIV-1) and murine leukemia virus (MLV) by blocking their entry into cells. Nevertheless, HIV-1 and MLV encode factors, Nef and glycosylated Gag respectively, that counteract SERINC3/5 *in vitro*. We recently developed SERINC3 and SERINC5 knockout mice to examine the *in vivo* function of these genes. We found that SERINC5 potently restricted retrovirus infection in a glycosylated Gag and envelope dependent manner. On the other hand, SERINC3 had no antiviral function. Our findings have implication in the development of therapeutics that target SERINC5 during retrovirus infection.

## INTRODUCTION

Cells have developed various restriction factors that counteract infection by inhibiting different points of the viral life cycle. Among these host restriction factors are the serine incorporator proteins (SERINC). The SERINC family of proteins consists of 5 members (SERINC1-5) and is conserved in all eukaryotes. They are all transmembrane proteins and are implicated with sphingolipid and phosphatidylserine biogenesis (1). Human SERINC3 and SERINC5 can inhibit a variety of retroviruses *in vitro*, including human immunodeficiency virus 1 (HIV-1) and murine leukemia virus (MLV) (2-5) with SERINC5 being more potently antiretroviral than SERINC3 (3, 4, 6). SERINC3 and SERINC5 are incorporated in budding virions (3, 4) and block the step of HIV envelope fusion and pore formation with the target cell membrane (7, 8).

MLV is divided into different subgroups on the basis of host range (9). Ecotropic MLVs, such as Friend MLV (F-MLV) infect solely mouse cells, amphotropic MLVs, such as 4070A, infect both human and mouse cells and xenotropic MLVs infect non murine cells (10). Glycosylated Gag (Glyco-Gag) is a viral protein expressed by both the ecotropic and amphotropic MLVs (11). Glyco-Gag is initiated from a CUG upstream of the polyprotein’s AUG, is highly glycosylated (12-15) and is cleaved to two fragments by an unknown protease (14, 16). The N’ terminal fragment is a transmembrane protein and the C’ terminal fragment is secreted from the infected cell (12, 17, 18). While Glyco-Gag is considered to be dispensable *in vitro*, glyco-Gag deficient viruses exhibit lower infectivity *in vivo* (19-21). Glyco-Gag is important *in vivo*, because it protects the virus from both the deleterious effects of mouse apolipoprotein B editing complex 3 (APOBEC3), a potent antiretroviral factor, and the induction of type I interferon response (11, 22-24). Glyco-Gag antagonizes the antiviral function of SERINC3 and SERINC5 *in vitro* by blocking the incorporation of SERINC proteins in the budding virions and leading to their lysosomal degradation (3, 4, 25, 26). Whether SERINC5 restricts retrovirus infection *in vivo* in a glyco-Gag dependent manner is currently unknown.

While a lot of work has been done to understand the role of SERINC proteins in retrovirus infection *in vitro*, nothing is known about the antiretroviral function of these proteins *in vivo*. A host factor that has antiretroviral function *in vitro* does not necessarily mean that it can restrict retrovirus infection *in vivo*. For example, SAMHD1, a potent retroviral factor, did not affect retrovirus infection in vivo (27). Here, for the first time, we examine the antiretroviral effect of SERINC5 *in vivo* and show that mouse SERINC5 (mSERINC5) restricts MLV infection *in vivo*. Moreover, SERINC5 restriction *in vivo* depends not only on the presence of glyco-Gag but also on the virus envelope. SERINC5 had no effect on F-MLV infectivity even when glyco-Gag was mutated; however, only when we substituted the F-MLV envelope with that of the amphotropic MLV 4070A envelope, we found that SERINC5 restricted MLV infection in a glyco-Gag dependent manner. Finally, unlike human SERINC3, mouse SERINC3 has no antiretroviral function both *in vivo* and *in vitro*.

## RESULTS

### Murine SERINC3 and SERINC5 are expressed in murine leukocytes and are not induced by F-MLV infection

To elucidate whether the mouse SERINC genes are expressed in leukocytes, cells that are naturally infected by F-MLV in mice (23, 28, 29), we isolated peripheral blood mononuclear cells (PBMCs) from 3 BL/6 mice (C57BL/6N-wild-type mice) and cell-sorted for T cells, B cells and dendritic cells (DCs). Transcription levels of SERINC genes were determined by RT-qPCR. We observed that only mSERINC1, −3 and −5 RNA were detected, while mSERINC2 and −4 were absent from these cell populations (Fig. 1A-C). To determine the effect of F-MLV infection on mSERINC3 and mSERINC5 transcription, we infected EL-4 (T lymphoblast cell line) and MutuDC1940 (a murine DC cell line (30)) with F-MLV wild-type (F-MLV WT) (0.1 MOI) followed by RT-qPCR to determine mSERINC3 and mSERINC5 transcription levels over time. No changes in the transcription levels of mSERINC5 and mSERINC3 were observed during infection (Fig. 1D, E). Thus, we concluded, that F-MLV infection does not affect mSERINC3 and mSERINC5 transcription.

**Fig. 1.**
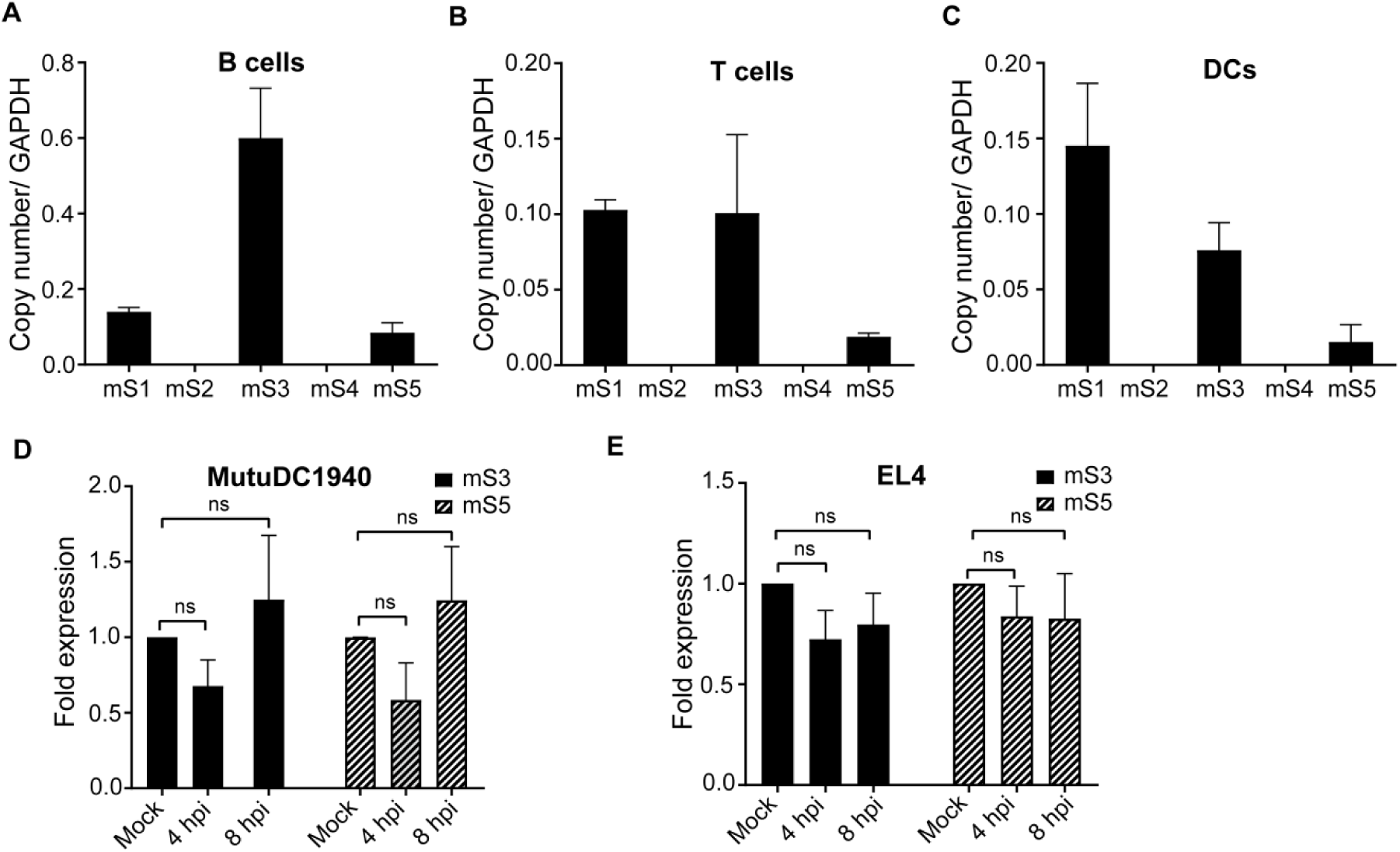
mSERINC3 and SERINC5 are constitutively expressed in murine leukocytes and are not induced by F-MLV infection. (A-C) mSERINC1-5 RNA copy number relative to GAPDH from (A) B cells (CD45R/B220+), (B) T cells (CD3+) and (C) Dendritic cells (DCs) (CD11c+) sorted from peripheral blood mononuclear cells isolated from the blood of C57BL/6 mice (n=3). (D-E) mSERINC3 and mSERINC5 transcripts fold expression relative to mock, normalized to GAPDH, in (D) MutuDC1940 and (E) EL4 cells, infected with F-MLV for 4h or 8h. Mock indicates mock-infected (PBS). Shown is the average of 3 independent experiments. Statistical significance was determined by one-way ANOVA and Tukey’s test. All results are presented as mean ± standard deviation (SD). ns, not significant. (mouse SERINC1-5, mS1-5; hour post infection, hpi)

### Generation of the SERINC5 -/- mice

As SERINC5 is the most potent antiretroviral member of the SERINC family of proteins (3-5), we generated SERINC5-deficient mice in the C57BL/6 background by targeting SERINC5 for deletion using clustered regularly interspersed short palindromic repeat (CRISPR)/ CRISPR associated 9 (Cas9) technology (Fig. 2A). Following electroporations to generate CRISPR knockout mice, we identified two founder lines and both were successful breeders. We isolated RNA from the tails of SERINC5+/+ (BL/6 mice), SERINC5+/- (heterozygous) and SERINC5-/- (homozygous knockout) mice and found that the SERINC5-/- mice had significantly lower SERINC5 transcripts than the SERINC5+/+, while the heterozygotes (+/-) had SERINC5 transcript levels intermediate between the SERINC5+/+ and the SERINC5-/- mice (Fig. 2B). Finally, fluorescence-activated cell sorting (FACS) analysis demonstrated that SERINC5-/- mice and BL/6 mice had similar levels of DCs, B cells, T cells and macrophages (Fig. 2C).

**Fig. 2.**
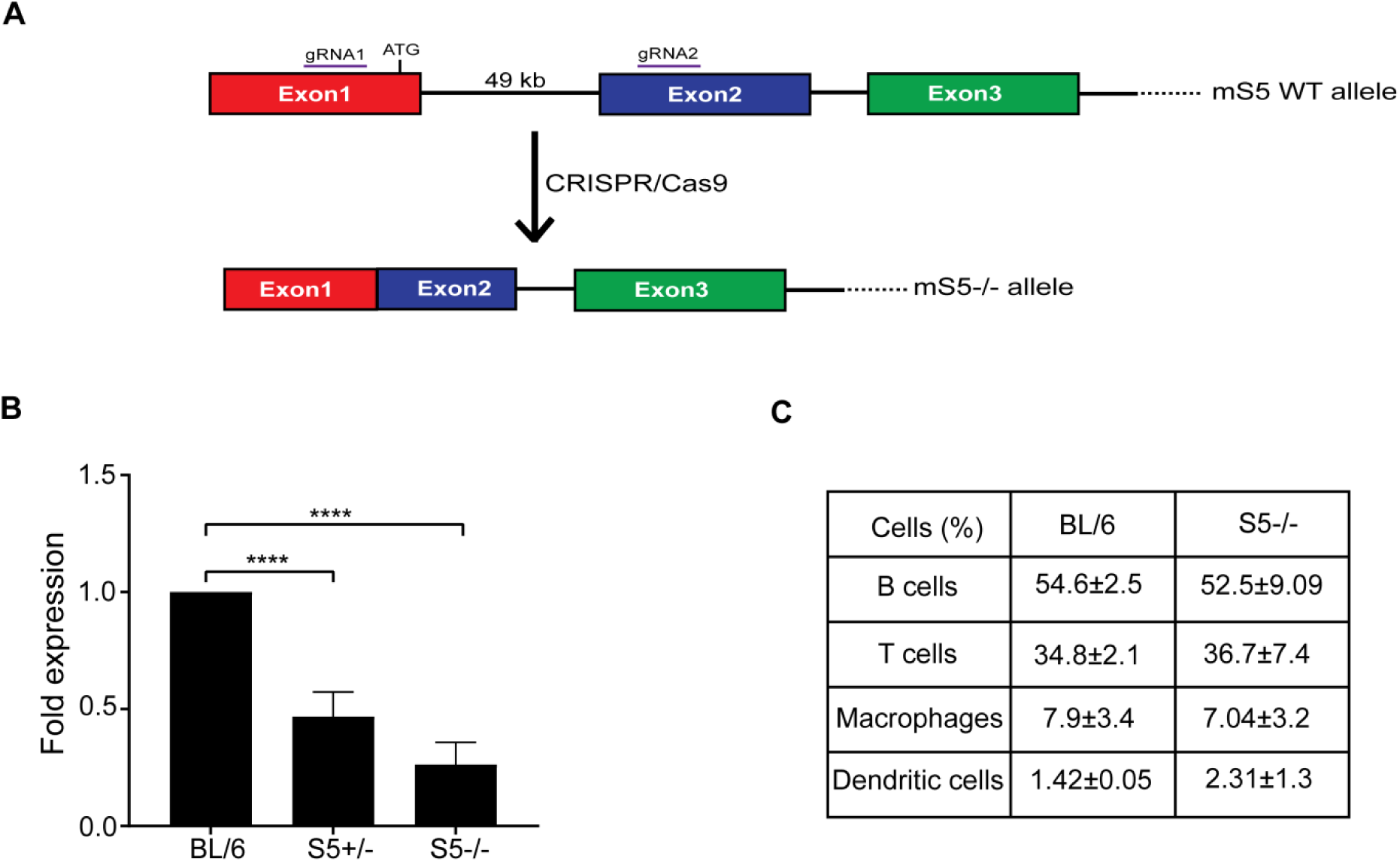
SERINC5-/- mice have lower levels of mSERINC5 and retain normal leukocytes populations. (A) Schematic of the SERINC5 allele in the SERINC5-/- mice. Two guide RNAs (gRNAs) were used to target exons 1 and 2. The genomic fragment between the two gRNAs was excised using CRISPR/Cas9 technology resulting in the abrogation of the SERINC5 gene. (B) Fold change of SERINC5 expression in the SERINC5-/- and SERINC5+/- mice relative to C57BL/6 mice. SERINC5 RNA levels in the SERINC5-/-, SERINC5+/-, and C57BL/6 mice were determined by RT-qPCR and normalized to β-actin. C57BL/6 (n = 6 mice), SERINC5+/- (n = 6 mice) and SERINC5-/- (n = 5 mice). (C) PBMCs from 4 C57BL/6 mice and 6 SERINC5-/- mice were stained with anti-CD45R/B220 (B cells), anti-CD3 (T cells), anti-F4/80 (Macrophages) and anti-CD11c (Dendritic cells) antibodies and analyzed by FACS. Values are presented as mean ± SD. There were no significant differences in the percentages of T, B, Macrophages and Dendritic cells among C57BL/6 and SERINC5-/- mice. For B, statistical significance was determined by one-way ANOVA and Tukey’s test; and unpaired t test for C, **, P <0.01, ****, P <0.0001. (SERINC5, S5; C57BL/6, BL/6)

### Murine SERINC3 and SERINC5 have no effect on ecotropic MLV infection *in vitro*

While the antiretroviral role of human SERINC3 and SERINC5 has been extensively examined (3-5), there is little information regarding the role of mSERINC3 and mSERINC5. To examine the effect of glyco-Gag on mSERINC3 and mSERINC5 during infection, we generated two F-MLV infectious clones with mutations in the glyco-Gag gene (Fig. 3A); one with two stop codons at amino acids 32 and 55 of glyco-Gag (gGag^-^F-MLV); previous work showed that introducing stop codons at these residues affected only glyco-Gag and not Gag translation (5). For the second virus, residues P31, Y36, L39 and R63 (gGag^mut^F-MLV), residues critical for the anti-SERINC5 function of glyco-Gag, were mutated to alanine (5, 25, 31). NIH3T3 cells were infected at MOI of 0.1 with gGag^-^F-MLV, gGag^mut^F-MLV and F-MLV WT and virus yields were determined at various times post infection by p30-CA ELISA while MLV DNA levels in the infected cells were measured by RT-qPCR. The gGag^-^F-MLV and the gGag^mut^F-MLV did not differ from the F-MLV WT in either p30 or MLV DNA levels (Fig. 3B, C), thus glyco-Gag is not essential for ecotropic MLV replication and infectivity *in vitro* (5, 20, 32, 33). Glyco-Gag blocks human SERINC5 and SERINC3 incorporation in nascent virions (3, 4, 34). To determine whether the glyco-Gag mutant viruses we generated affect the incorporation of mSERINC3 and mSERINC5 in budding MLV particles, we co-transfected 293T cells with either F-MLV WT or the F-MLV constructs with mutations in the glyco-Gag gene and with either mSERINC3 or mSERINC5. Virus released was examined for mSERINC3 and mSERINC5 content by western blots. We found that gGag^-^F-MLV and gGag^mut^F-MLV particles had higher levels of mSERINC3 and mSERINC5 incorporated in their virions when compared to F-MLV WT particles (Fig. 3D). Thus, we concluded that glyco-Gag blocks mSERINC3 and mSERINC5 incorporation in the nascent virions.

**Fig. 3.**
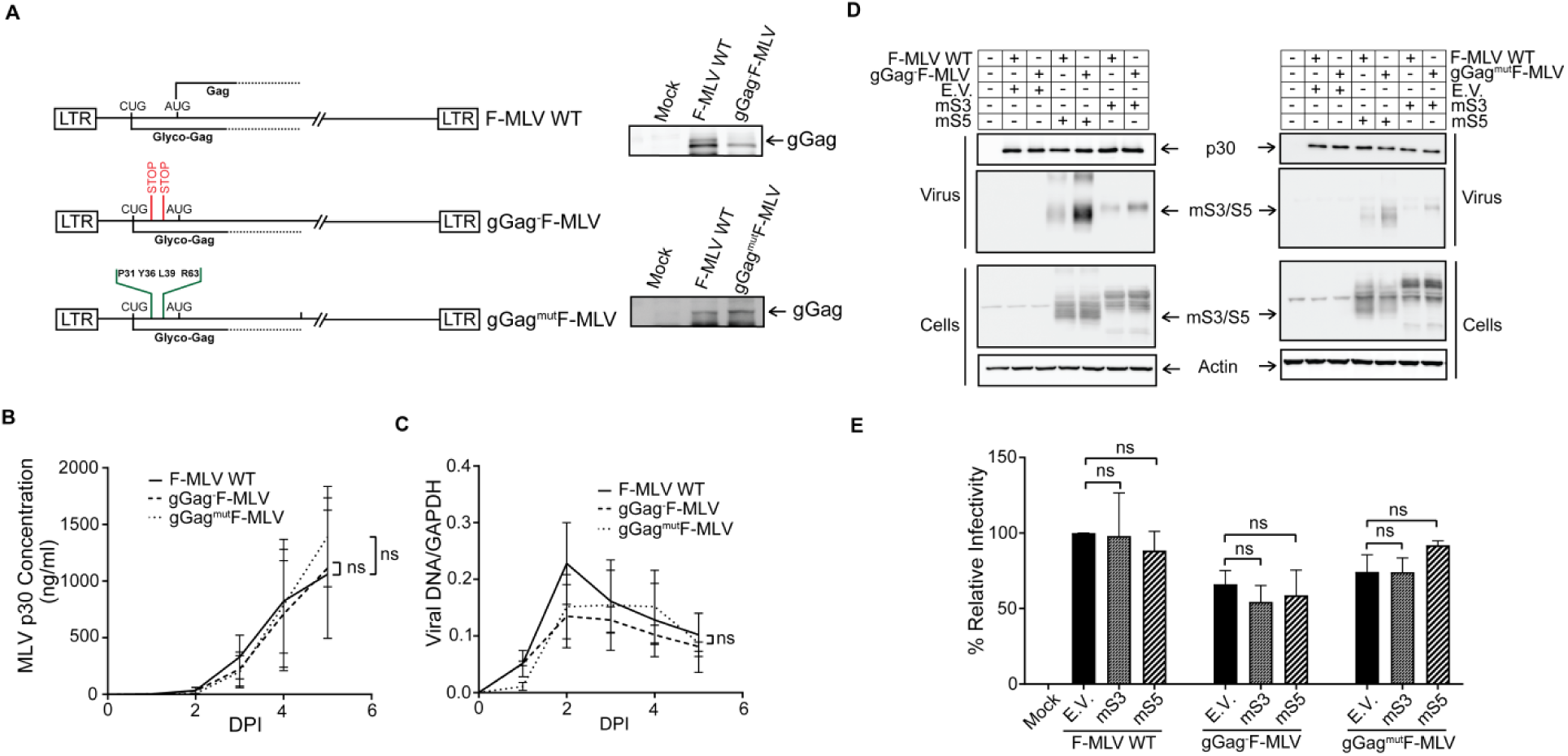
mSERINC3 and mSERINC5 have no effect on ecotropic MLV infection *in vitro*. (A) Schematic diagram of glyco-Gag (gGag) mutant MLV constructs. Positions of translation start codons (CUG for glyco-Gag and AUG for Gag) and the two stop codons inserted at amino acids 32 and 55 within glyco-Gag reading frame to generate gGag^-^F-MLV are shown. In the case of gGag^mut^F-MLV construct, glyco-Gag residues P31, Y36, L39 and R63 were mutated to alanine (A). Shown on the right are immunoblots of lysates from 293T cells transfected with F-MLV WT, gGag^-^F-MLV and gGag^mut^F-MLV constructs and probed with a goat anti-MLV antibody for the detection of gGag. (B-C) Growth curves of the gGag mutant viruses. NIH3T3 cells were infected with F-MLV WT, gGag^-^F-MLV and gGag^mut^F-MLV (MOI 0.1). Virus replication was determined (B) by performing p30 (CA) ELISA assays to determine virus levels in the culture supernatants at the indicated time points and (C) by RT-qPCR for MLV DNA levels normalized to GAPDH in the infected NIH3T3. (D) mSERINC3 and mSERINC5 incorporation in the budding virions is glyco-Gag dependent. 293T cells were co-transfected with F-MLV WT/ gGag^-^ F-MLV/ gGag^mut^F-MLV and either mSERINC3, mSERINC5 or empty vector as indicated. Forty eight hours post transfection, cells and released virus in the culture media were harvested and the indicated proteins were analyzed by immunoblotting using anti-MLV p30, anti-HA (for detection of mSERINC3 and mSERINC5) and anti-β-actin antibodies. (E) mSERINC3 and mSERINC5 do not affect F-MLV infectivity *in vitro*. Mus Dunni cells were infected with equal amounts of F-MLV WT or gGag^-^F-MLV or gGag^mut^F-MLV virus produced in the presence of mSERINC3, mSERINC5 or empty vector. Genomic DNA was isolated 5 h post infection (hpi) and MLV DNA levels were measured by RT-qPCR. Viral DNA levels normalized to GAPDH were used to calculate the percentage (%) of relative infectivity with respect to F-MLV WT virus produced in the presence of empty vector. All results are presented as mean ± SD. Statistical significance was determined by unpaired t test (two-tailed) for data points at 5 DPI (B, C) and by one-way ANOVA and Tukey’s test (E). ns, not significant. Results are shown for n = 3 independent experiments (A-E); representative immunoblotting results are shown in (A, D). (mouse SERINC3, mS3; mouse SERINC5, mS5; empty vector, E.V.; Glyco-Gag, gGag)

We next examined the effect of mSERINC3 and mSERINC5 on F-MLV infection. Previous work has shown that although hSERINC3 and hSERINC5 do not affect virus release from producer cells, they blocked MLV infectivity in a glyco-Gag dependent manner in target cells (3-5, 25, 34). To determine whether mSERINC3 and mSERINC5 packaged in the MLV particles blocked MLV infection, we infected Mus Dunni cells with equal amounts of F-MLV WT, gGag^-^F-MLV and gGag^mut^F-MLV generated in the presence of either mSERINC3 or mSERINC5. DNA was isolated from the infected cells and viral DNA levels were measured by RT-qPCR. Interestingly, F-MLV infectivity was unaffected regardless of the levels of mSERINC3 or mSERINC5 incorporated in the nascent virions (Fig. 3E), demonstrating that while mSERINC3 and mSERINC5 were excluded from F-MLV in a glyco-Gag dependent manner, these two host factors had no effect on F-MLV infectivity.

### Murine SERINC5 does not restrict ecotropic MLV infection *in vivo*

While mSERINC3 and mSERINC5 did not affect F-MLV infectivity when glyco-Gag was mutated, we speculated that it is possible that *in vivo* we may observe something different. *In vitro* data do not necessarily reflect what happens *in vivo.* During *in vivo* infections MLV infects a variety of cells, and the immune system, due to its activation by MLV infection, might result in changes in the levels of host genes like SERINC5 that might affect the outcome of infection. To determine the role of SERINC5 on retrovirus infection *in vivo*, we infected the SERINC5-/- mice we developed with F-MLV WT, gGag^-^F-MLV and gGag^mut^F-MLV, and 10 days post infection (dpi) virus titers in their spleens were measured. We found that both BL/6 (SERINC5+/+) and SERINC5-/- mice infected with F-MLV WT had similar virus titers in their spleens at 10 dpi (Fig. 4). Interestingly, gGag^-^F-MLV and gGag^mut^F-MLV also replicated to similar levels in the spleens of both BL/6 and SERINC5-/- mice, albeit both viruses grew at significantly lower levels when compared to F-MLV WT (Fig. 4).

**Fig. 4.**
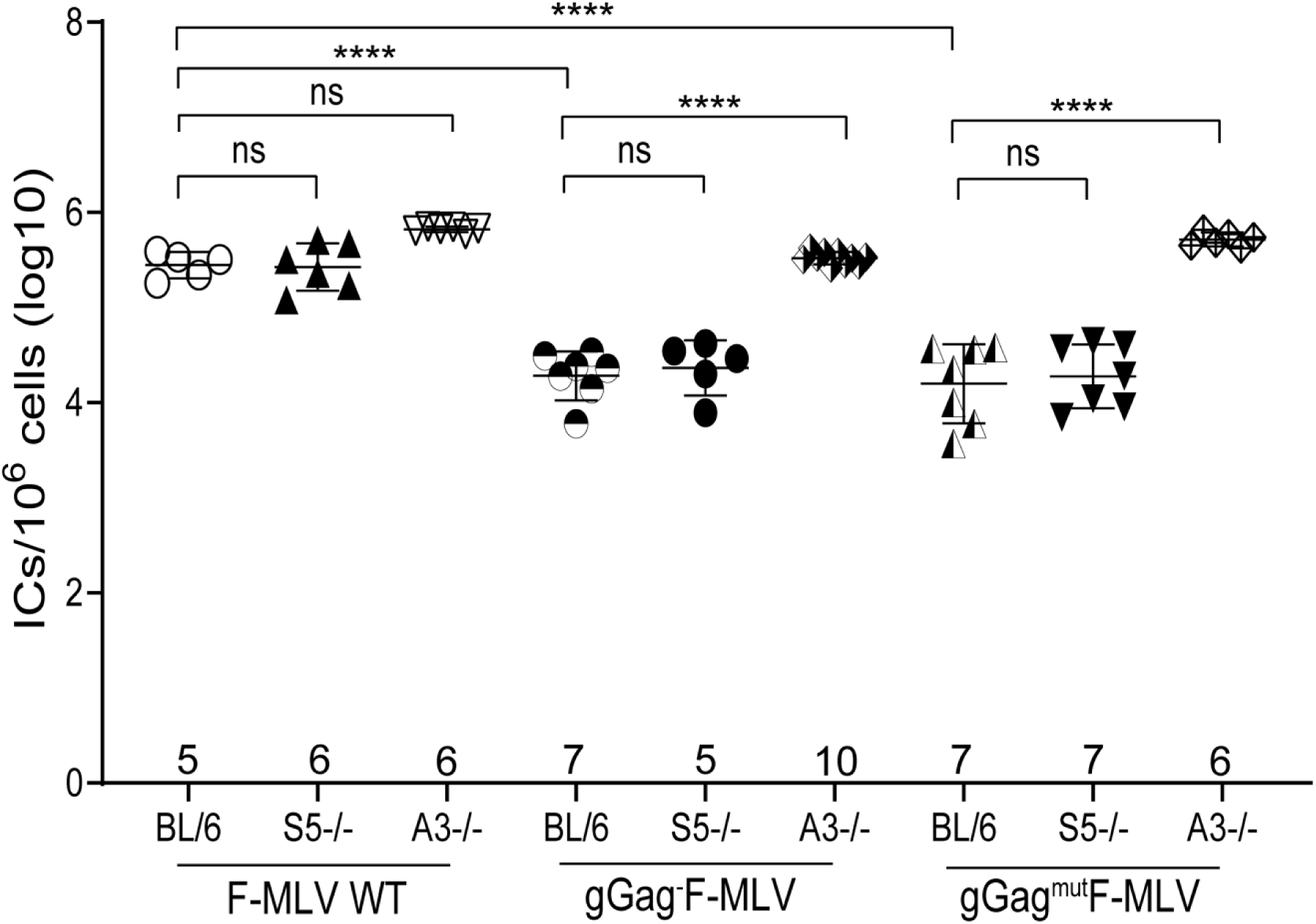
SERINC5 has no effect on ecotropic MLV infection *in vivo*. Newborn mice were infected with F-MLV WT, gGag^-^F-MLV and gGag^mut^F-MLV virus and virus titers in the spleens were measured 10 days post infection. Each point represents the titer obtained from an individual mouse and the average for each group is shown by a horizontal line. Mice were derived from 2-3 litters per genotype; the C57BL/6 and SERINC5-/- mice are littermates. Numbers of mice used in each group are indicated in the X-axis. All results are presented as mean ± SD. Statistical significance was determined by one-way ANOVA and Tukey’s test. ****, p <0.0001; ns, not significant. (C57BL/6, BL/6; APOBEC3, A3; Glyco-Gag, gGag; Infectious centers, ICs)

As a positive control to our *in vivo* experiments, we used APOBEC3-/- (A3-/-) mice, where a glyco-Gag mutant MLV has been shown to replicate similarly to wild-type MLV in the spleens of the infected mice (11, 22, 24). We infected APOBEC3-/- mice with F-MLV WT, gGag^-^F-MLV and gGag^mut^F-MLV and similar to what has been previously reported (11, 22, 24), all three viruses replicated to similar levels in the spleens of the infected APOBEC-/- mice (Fig. 4).

Thus, we concluded that in the context of the ecotropic F-MLV, SERINC5, unlike APOBEC3, exerts no *in vivo* antiviral function regardless of the presence of glyco-Gag.

### The amphotropic MLV envelope renders MLV vulnerable to SERINC5 restriction in a glyco-Gag dependent manner *in vitro*

It was previously shown that envelopes from different HIV subtypes could overcome SERINC5 restriction without preventing SERINC5 incorporation in the virions (35, 36). Therefore, virus susceptibility to human SERINC3 and SERINC5 restriction is not only dependent on glyco-Gag, but also on the viral envelope (2-5, 35). We thus hypothesized that the envelope of MLV might play a crucial role in mSERINC5-mediated restriction *in vivo*. Amphotropic MLV envelope, just like xenotropic MLV envelope, rendered MLV pseudoviruses sensitive to SERINC5 restriction (5, 33) and the glyco-Gag sequences of amphotropic and ecotropic MLV are 90% identical (data not shown) when compared. We developed chimeric F-MLV constructs with the aforementioned mutations in glyco-Gag but the envelope was replaced with that of the amphotropic MLV strain 4070A (Fig. 5A) - gGag^-^F-MLV/Ampho^Env^ and gGag^mut^F-MLV/Ampho^Env^. As a control we also replaced the envelope of F-MLV WT with that of the amphotropic 4070A-F-MLV/Ampho^Env^ (Fig. 5A). To determine whether glyco-Gag still affected mSERINC3 and mSERINC5 incorporation in virions with the amphotropic envelope, we co-transfected 293T cells with the chimeric envelope viral constructs and either mSERINC3 or mSERINC5. Western blots on purified virions showed that mSERINC3 and mSERINC5 were incorporated in a glyco-Gag dependent manner; virions where glyco-Gag was either absent or mutated (gGag^-^F-MLV/Ampho^Env^ and gGag^mut^F-MLV/Ampho^Env^) had higher mSERINC3 and mSERINC5 levels packaged than virions with intact glyco-Gag (F-MLV/Ampho^Env^) (Fig. 5B). Thus, we concluded glyco-Gag blocks the incorporation of mSERINC3 and mSERINC5 in the viral particles independently of the virus envelope.

**Fig. 5.**
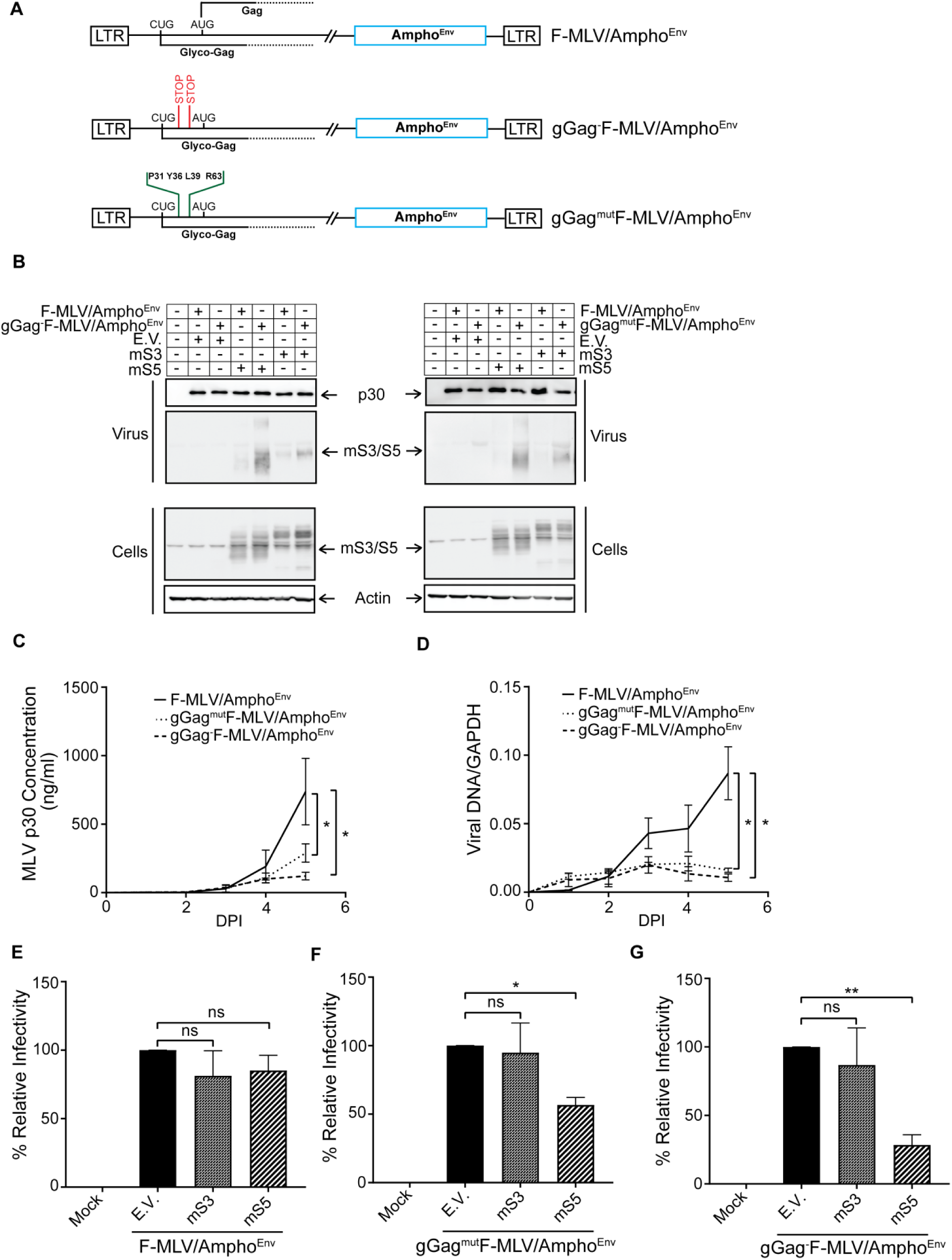
Amphotropic envelope renders MLV sensitive to mSERINC5 restriction *in vitro*. (A) Schematic of the chimeric F-MLV constructs expressing the amphotropic envelope. (B) Mouse SERINC3 and mouse SERINC5 incorporation in the budding virions is dependent on glyco-Gag and not on the viral envelope. 293T cells were co-transfected with F-MLV/Ampho^Env^, gGag^-^F-MLV/Ampho^Env^ and gGag^mut^F-MLV/Ampho^Env^ and either mSERINC3 or mSERINC5 or empty vector as indicated. Forty eight hours post transfection, cells and released virus in the culture media were harvested and analyzed by immunoblotting using anti-MLV p30, anti-HA (for detection of mSERINC3 and mSERINC5) and anti-β-actin antibodies. (C-D) Mutations in glyco-Gag result in loss of infectivity in the chimeric viruses. NIH3T3 cells were infected with F-MLV/Ampho^Env^, gGag^-^F-MLV/Ampho^Env^ and gGag^mut^F-MLV/Ampho^Env^ (MOI 0.1). Virus replication was monitored (C) in the culture supernatants by MLV p30 CA ELISA assays at the indicated time points, and (D) in the infected cells by isolating genomic DNA and determining by RT-qPCR MLV DNA levels followed by normalization to GAPDH. (E-G) mSERINC3 and mSERINC5 restrict virions with amphotropic envelope *in vitro*. Mus Dunni cells were infected with equal amounts of F-MLV/Ampho^Env^ (E), gGag^mut^F-MLV/Ampho^Env^ (F) and gGag^-^F-MLV/Ampho^Env^ (G) produced in the presence of either mSERINC3 or mSERINC5 or empty vector. Cells were harvested 5 hours post infection and MLV DNA levels were measured by RT-qPCR and normalized to GAPDH. Percentage (%) of relative infectivity was determined with respect to virus generated in the presence of empty vector. All results are presented as mean ± SD. Statistical significance was determined by unpaired t test (two-tailed) for data points at 5 DPI (C, D) and by one-way ANOVA and Tukey’s test (E-G). *, p < 0.05; **, p < 0.01; ns, not significant. Results are shown for n = 3 independent experiments (B-G); representative immunoblotting results are shown in B. (mouse SERINC3, mS3; mouse SERINC5, mS5; empty vector, E.V.)

We then performed growth curves on the different chimeric viruses we generated. Mus Dunni cells were infected at MOI of 0.1 and virus levels were determined at various times post infection by p30-CA ELISA and by RT-qPCR measuring MLV DNA levels in the infected cells. Remarkably, unlike what we saw with the viruses that had ecotropic MLV envelope, the glyco-Gag mutant chimeric viruses carrying amphotropic MLV envelope (gGag^-^F-MLV/Ampho^Env^ and gGag^mut^F-MLV/Ampho^Env^) replicated considerably less when compared to the chimeric virus carrying an intact glyco-Gag (F-MLV/Ampho^Env^) (Fig. 5C, D). Therefore, due to the different growth kinetics of the three viruses (gGag^-^F-MLV/Ampho^Env^, gGag^mut^F-MLV/Ampho^Env^, and F-MLV/Ampho^Env^) we have abstained from making any direct comparisons among the different viruses. To examine the effect of mSERINC3 and mSERINC5 on the infectivity of the chimeric viruses, we infected Mus Dunni cells with equal amounts of the three chimeric viruses (gGag^-^F-MLV/Ampho^Env^, gGag^mut^F-MLV/Ampho^Env^, and F-MLV/Ampho^Env^) that were generated in the presence of either mSERINC3 or mSERINC5. We then measured MLV DNA levels by RT-qPCR and found that the glyco-Gag expressing chimeric virus, F-MLV/Ampho^Env^, replicated the same in infected Mus Dunni cells regardless of the presence of mSERINC3 or mSERINC5 (Fig. 5E). On the other hand, chimeric viruses, in which glyco-Gag was either mutated or deleted (gGag^mut^F-MLV/Ampho^Env^ and gGag^-^F-MLV/Ampho^Env^), replicated very poorly in the presence of mSERINC5 (Fig. 5F, G). Interestingly, incorporation of mSERINC3 in the chimeric virions, even when glyco-Gag was disrupted, had no effect on virus infectivity (Fig. 5F, G). This is particularly interesting as the human homologue of mSERINC3, human SERINC3, has antiretroviral functions (3, 4, 37). Thus, we concluded that mSERINC5, similar to human SERINC5, restricts retrovirus infection *in vitro* in a glyco-Gag and envelope dependent manner, while mSERINC3 has no antiretroviral function *in vitro*.

### Murine SERINC5 restricts MLV infection *in vivo* in a glyco-Gag and envelope dependent manner

The change in the susceptibility to mSERINC5 restriction upon replacing the envelope of the ecotropic F-MLV to that of the amphotropic MLV, 4070A, provides further proof that the envelope of the viral particles is critical for SERINC5 restriction. To examine if the viral envelope affects SERINC5 restriction *in vivo*, we used the chimeric viruses F-MLV/Ampho^Env^, gGag^-^F-MLV/Ampho^Env^, and gGag^mut^F-MLV/Ampho^Env^ to infect SERINC5-/- and BL/6 mice. Titers in the spleens of the infected mice were measured 10 dpi. In the case of F-MLV/Ampho^Env^, both SERINC5-/- and BL/6 mice had similar viral titers in their spleens, which is attributed to the fact that the virus has an intact glyco-Gag (Fig. 6A). Interestingly, in the case of the gGag^-^F-MLV/Ampho^Env^, infection was abrogated for all animals independent of genotype (SERINC5-/-, A3-/- and BL/6 mice) (Fig. S1). This was quite surprising, as in the context of the ecotropic envelope, deletion of glyco-Gag only resulted in reduced virus titers in the spleens of the infected BL/6 and SERINC5-/- mice (Fig. 4) and not abrogation of infection as we see with the amphotropic envelope containing chimeric virus. Remarkably, in the case of gGag^mut^F-MLV/Ampho^Env^, we observed that virus titers in the spleens of the SERINC5-/- mice were significantly higher than those observed in the BL/6 mice (Fig. 6B). This finding not only demonstrates that mSERINC5 restricts MLV infection *in vivo* but also that MLV susceptibility to SERINC5 inhibition is dependent on both the viral envelope and the presence of glyco-Gag.

**Fig. 6.**
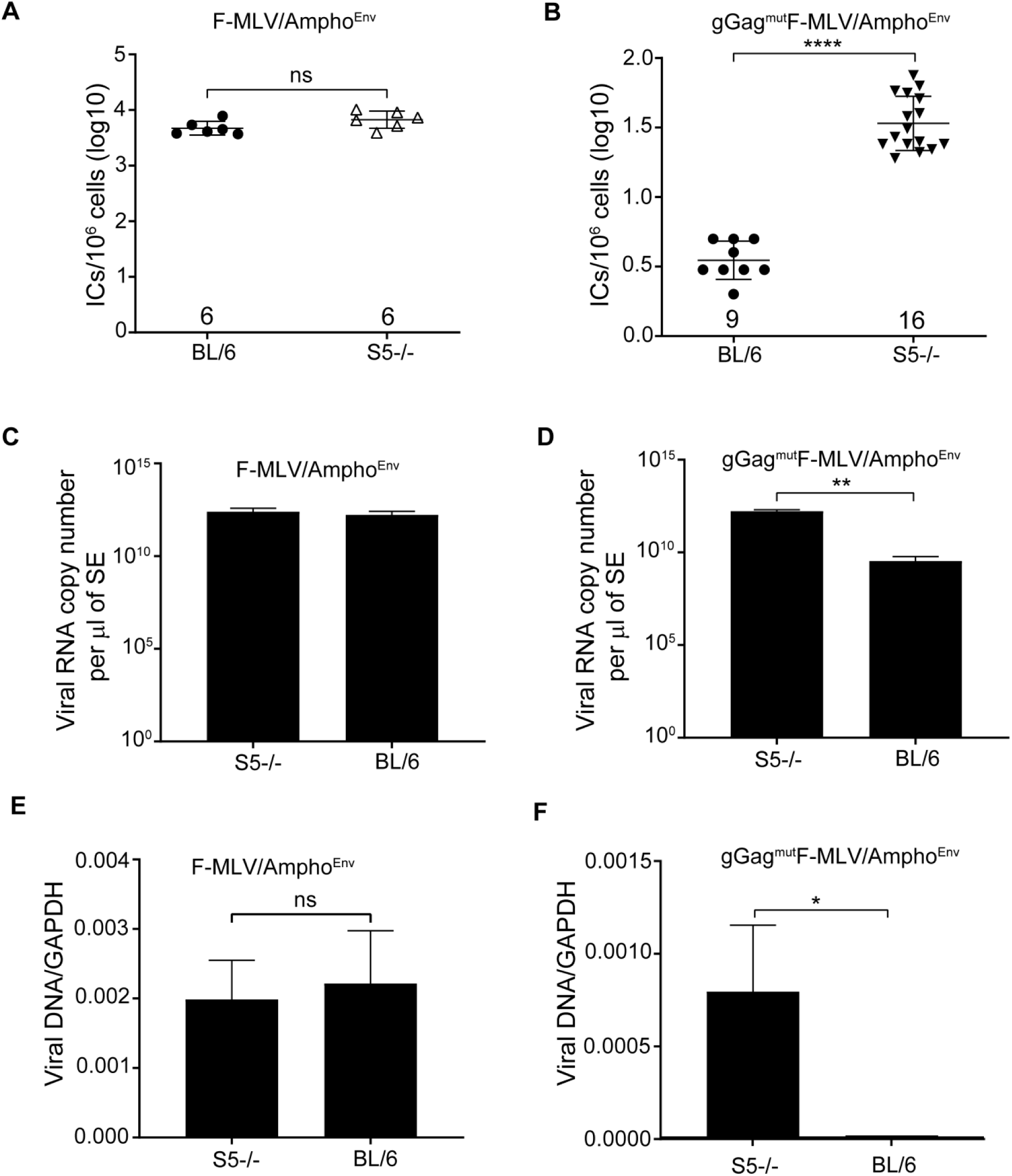
mSERINC5 restricts MLV infection *in* vivo in an envelope and glyco-Gag dependent manner. (A, B) Newborn mice were infected with F-MLV/Ampho^Env^ (A) and gGag^mut^F-MLV/Ampho^Env^ (B). Virus titers in spleens were measured 10 days post infection. Each point represents the titer obtained from an individual mouse and the mean for each group is shown by a horizontal line. Mice were derived from 4-6 litters each; the C57BL/6 and SERINC5-/- are the littermates. Numbers of mice used in each group are indicated in the X-axis. (C, D) Newborn mice were infected with (C) F-MLV/Ampho^Env^ or (D) with gGag^mut^F-MLV/Ampho^Env^ and splenic extracts were prepared 16 days post infection. Viral RNA copy number was measured in the splenic extracts using RT-qPCR. For (C) n = 4 (C57BL/6); n = 8 (SERINC5-/-) and for (D) n = 3 (C57BL/6); n = 3 (SERINC5-/-). (E, F) Equal amounts of viral RNA copies (from C and D) were used to infect Mus Dunni cells. Genomic DNA was isolated 19 h post infection and viral DNA levels were measured by RT-qPCR and normalized to GAPDH are presented. All results are presented as mean ± SD. Statistical significance was determined by Mann-Whitney test (two-tailed) (A, B) and by unpaired t test (C, D). *, < 0.05; **, < 0.01; ****, <0.0001; ns, not significant. (SERINC5, S5; BL/6, C57BL/6; SE, splenic extracts; Infectious centers, ICs)

Our *in vitro* data show that glyco-Gag mutant viral particles with the amphotropic envelope (gGag^mut^F-MLV/Ampho^Env^) have decreased infectivity when mSERINC5 is packaged in the virions. The spleen is one of the major organs that becomes infected by MLV *in vivo* (11, 38-41) Thus, viral particles purified from the spleens of infected BL/6 mice should have reduced infectivity when compared to those derived from SERINC5-/- infected mice. To test this, we purified virions from the spleens of BL/6 and SERINC5-/- mice infected with either gGag^mut^F-MLV/Ampho^Env^ or F-MLV/Ampho^Env^. We initially measured viral RNA levels in the splenic extracts and found that F-MLV/Ampho^Env^ viral RNA levels were similar in both BL/6 and SERINC5-/- mice (Fig. 6C), while gGag^mut^F-MLV/Ampho^Env^ viral RNA levels were higher in the splenic extracts of the SERINC5-/- mice compared to BL/6 mice (Fig. 6D), which reflects the higher infection levels of the former (Fig. 6B). Subsequently, we infected Mus Dunni cells with equal amounts of gGag^mut^F-MLV/Ampho^Env^ or F-MLV/Ampho^Env^ viral particles isolated from BL/6 or SERINC5-/- mice after normalizing to viral RNA levels. We measured MLV DNA levels by RT-qPCR and found no difference in the infectivity of F-MLV/Ampho^Env^ virions isolated from either BL/6 or SERINC5-/- mice (Fig. 6E). On the other hand, gGag^mut^F-MLV/Ampho^Env^ virons isolated from SERINC5-/- mice were more infectious than those from BL/6 mice (Fig. 6F). The absence of an antibody that could detect endogenous levels of mSERINC5 prevented us from examining mSERINC5 incorporation in the nascent virions *in vivo*. Taken together, these data suggest that mSERINC5 is packaged into virions isolated from the spleens of infected mice in a glyco-Gag dependent manner and limits viral infectivity.

### SERINC5 and APOBEC3 synergistically act to restrict retrovirus infection *in vivo*

Mouse APOBEC3 potently restricts MLV infection *in vivo* by blocking reverse transcription (11, 22, 24, 38, 41, 42). SERINC5 blocks virus-cell fusion (7, 8). As mouse APOBEC3 and mSERINC5 target different steps of the retrovirus life cycle and both restrict MLV infection *in vivo*, we asked whether these two factors act synergistically to block retrovirus infection *in vivo*. To examine any synergistic effect between mouse APOBEC3 and mSERINC5, we developed double knockout mice for these two genes, APOBEC3/SERINC5-/- mice. We infected APOBEC3/SERINC5-/- mice with F-MLV/Ampho^Env^ and gGag^mut^F-MLV/Ampho^Env^ and 10 dpi we harvested their spleens to determine viral titers. F-MLV/Ampho^Env^ replicated at similar levels in the spleens of both APOBEC3-/- and APOBEC3/SERINC5-/- mice (Fig. 7A). In the case of gGag^mut^F-MLV/Ampho^Env^, the APOBEC3/SERINC5-/- mice had higher viral titers in their spleens than either the APOBEC3-/- mice or the SERINC5-/- mice (Fig. 7B). Therefore, we concluded that mouse APOBEC3 and mSERINC5 act synergistically to block retrovirus infection *in vivo*.

**Fig. 7.**
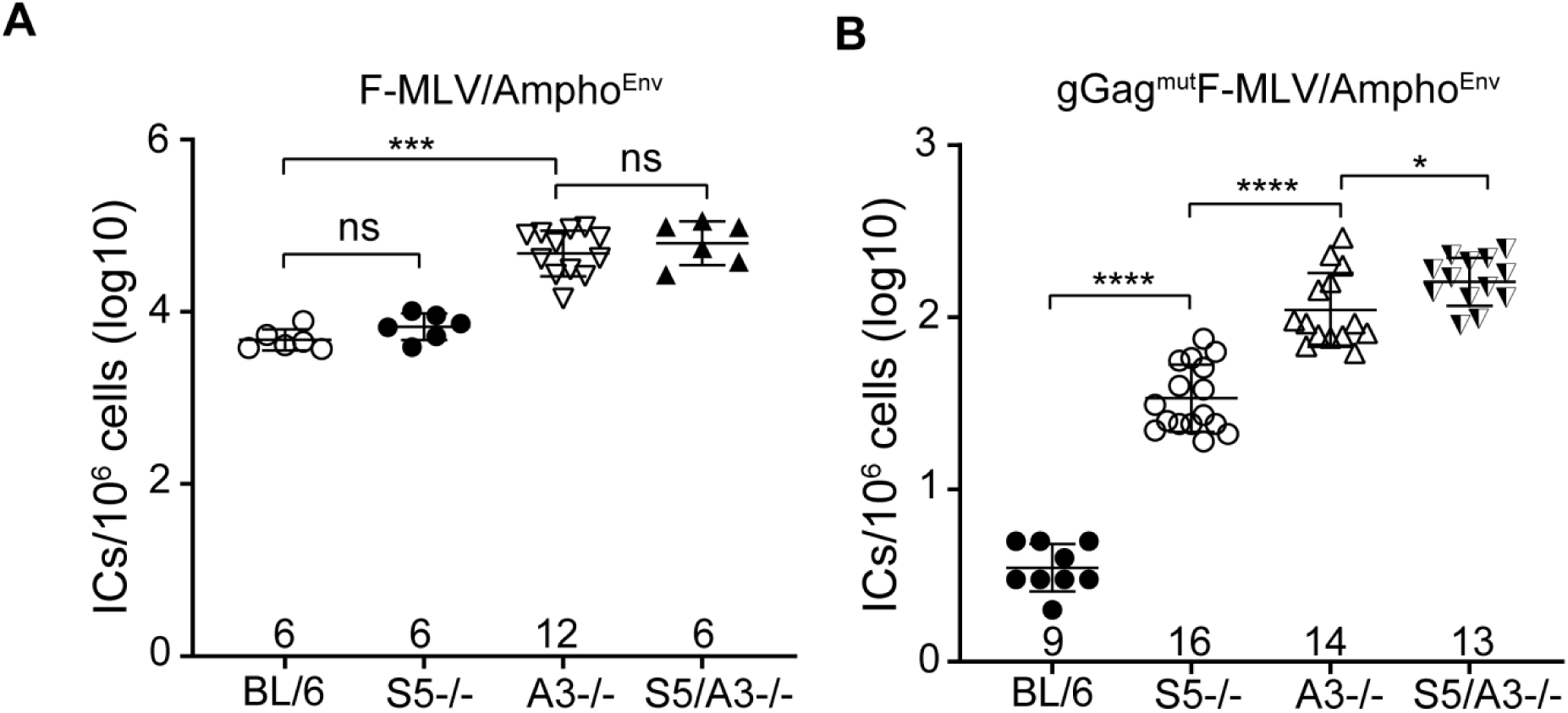
SERINC5 and APOBEC3 have synergistic effect on MLV infection *in vivo.* Newborn mice were infected with F-MLV/Ampho^Env^ (A) and gGag^mut^F-MLV/Ampho^Env^ (B). Virus titers in the spleens were measured 10 days post infection. Each point represents the titer obtained from an individual mouse and the mean for each group is shown by a horizontal line. Mice were derived from 2-6 litters each; the BL/6 and SERINC5 are the littermates. Numbers of mice used in each group are indicated in the X-axis. All results are presented as mean ± SD. Virus titers for BL/6 and SERINC5-/- mice shown in A and B for F-MLV/Ampho^Env^ and gGag^mut^F-MLV/Ampho^Env^ are duplicated from Fig. 6A and B respectively. Statistical significance was determined by Mann-Whitney test (two-tailed). *, <0.5; ***, p = 0.0001 ****, <0.0001; ns, not significant. (SERINC5, S5; C57BL/6, BL/6; APOBEC3, A3)

### Murine SERINC3 has no effect on MLV restriction *in vivo*

SERINC3 is another member of the SERINC protein family that has antiretroviral functions (3, 4). To determine whether mSERINC3 could exert an antiretroviral function *in vivo*, we acquired mice from the Wellcome Trust Sanger Institute Mouse Genetics Project with loxP sites flanking exon 3 of SERINC3. We crossed these mice with CMV-Cre expressing mice (Cre Deleter,Taconic) to develop complete -/- mice for mSERINC3 (Fig. 8A), as deletion of exon 3 results in a truncated mSERINC3 RNA transcript with a frameshift mutation that leads to a premature stop codon. Because there is no antibody to detect endogenous mSERINC3 protein levels, we probed for mSERINC3 mRNA levels by RT-qPCR and found that the SERINC3-/- mice had significantly lower levels of mSERINC3 transcripts when compared to BL/6 mice (Fig. 8B). SERINC3-/- mice had no apparent physical or behavioral deficits and similar leukocyte populations as BL/6 mice (Fig. 8C). Subsequently, we infected SERINC3-/- mice with either gGag^mut^F-MLV/Ampho^Env^ or F-MLV/Ampho^Env^. At 10 dpi, both viruses replicated the same in the spleens of either BL/6 or mSERINC3-/- mice (Fig. 8D, E). Thus, mSERINC3, unlike human SERINC3, has no effect on MLV infection *in vivo*.

**Fig. 8.**
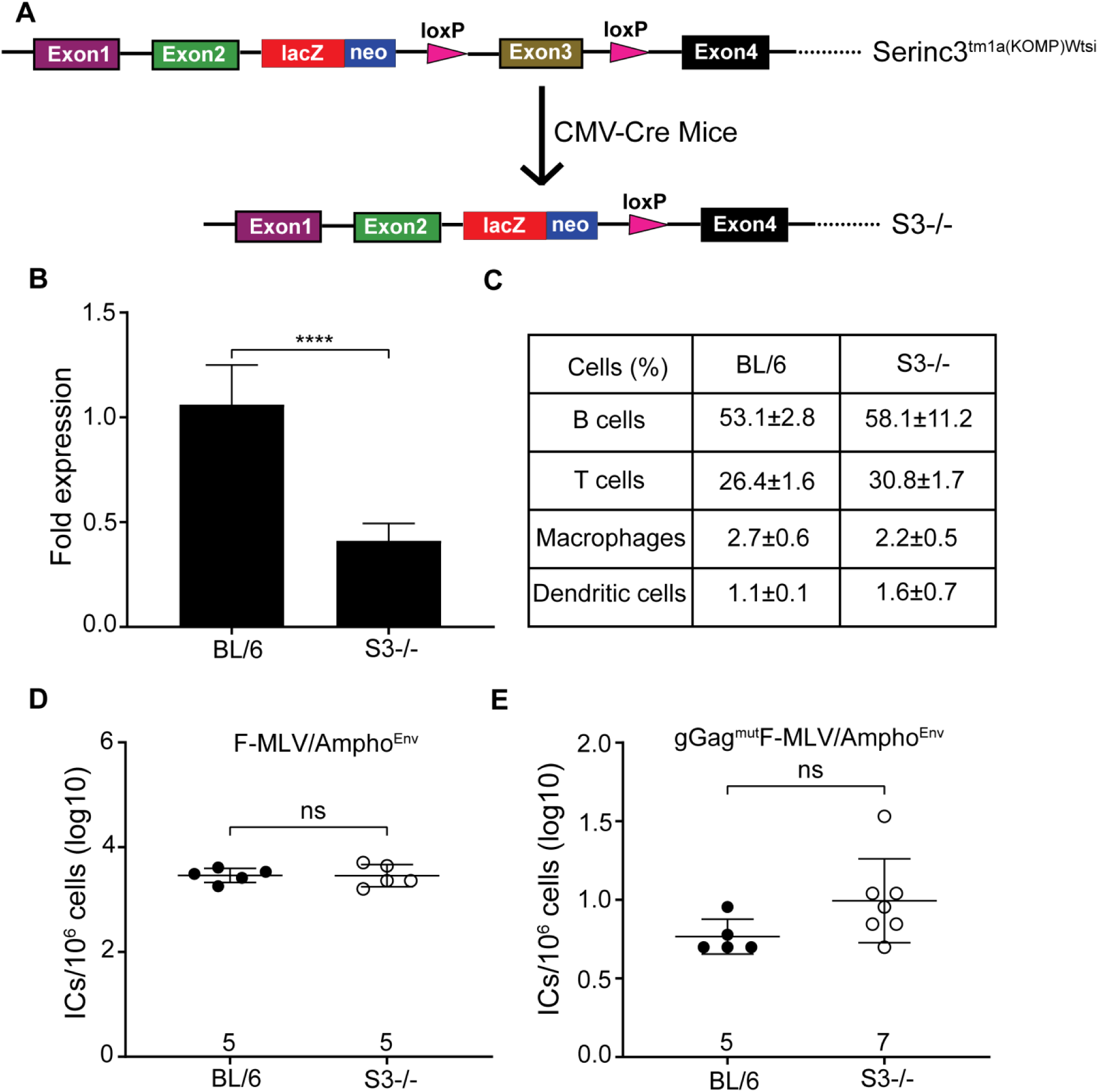
mSERINC3 has no effect on MLV infection *in vivo*. (A) Schematic of the derivation of SERINC3-/- mice. Serinc3^tm1a(KOMP)Wtsi^ mice provided by the Wellcome Trust Sanger Institute were crossed with the Cre-deleter mice (Taconic) leading to the loss of exon 3 and a frameshift mutation abrogating the SERINC3 gene. (B) Total RNA was isolated from C57BL/6 (n = 9) and SERINC3-/- (n = 9) mice and SERINC3 levels were measured by RT-qPCR and normalized to GAPDH. SERINC3 transcript fold expression was determined relative to that in C57BL/6 mice. (C) PBMCs from 3 C57BL6 mice and 4 SERINC3-/- mice were stained with anti-CD45R/B220 (B cells), anti-CD3 (T cells), anti-F4/80 (Macrophages) and anti-CD11c (Dendritic cells) and analyzed by FACS. Values are presented as mean ± SD. There were no significant differences in the percentages of T, B, Macrophages and Dendritic cells among C57BL/6 and SERINC3-/- mice. (D, E) C57BL/6 and SERINC3-/- newborn mice were infected with F-MLV/Ampho^Env^ (D) and gGag^mut^F-MLV/Ampho^Env^ (E). Virus titers in spleens were measured 10 days post infection. Each point represents the titer obtained from an individual mouse and the mean for each group is shown by a horizontal line. Mice were derived from 2-4 litters each. Numbers of mice used in each group are indicated in the X-axis. All results are presented as mean ± SD. Statistical significance was determined by unpaired t test (B, C) and by Mann-Whitney test (two-tailed) (D, E). ****, < 0.0001; ns, not significant. (SERINC3, S3; C57BL/6, BL/6; Infectious centers, ICs)

### MLV sensitivity to mSERINC5 is independent of the route of virus entry into target cells

Previous studies on the envelope of ecotropic MLVs like Moloney MLV (MoMLV) and F-MLV, have shown that virus internalization varies with the host cell (43-45). Our studies show that infection of NIH3T3 and Mus Dunni cells, both murine fibroblasts, was affected by the presence of ammonium chloride, a pH inhibitor, (Fig. S2A), which, in agreement with previous reports, suggested that virus fusion takes place in a low pH-environment after receptor-mediated endocytosis (44-46). On the other hand, in accord with previous reports (44-49), ecotropic MLV infection is pH independent in XC cells, a rat sarcoma cell line, as we did not observe any effect on virus infection, when ammonium chloride was added to the cells (Fig. S2A). In the case of amphotropic MLV, fusion happens on the cell surface and occurs at extracellular neutral pH (45, 48). However, another study found that entry of amphotropic MLV into NIH 3T3 cells was pH dependent (47). Similar to that report, we found that treatment of NIH 3T3 cells and Mus Dunni with ammonium chloride blocked both ecotropic and amphotropic MLV infection (Fig. S2B). To rule out whether the route of entry of ecotropic MLVs is responsible for the resistance of F-MLV to mSERINC5 restriction, we infected XC cells (where entry is by surface fusion and pH independent) with equal amounts of F-MLV WT, gGag^mut^F-MLV, gGag^mut^F-MLV/Ampho^Env^ or F-MLV/Ampho^Env^ generated in the presence of either mSERINC5 or empty vector. XC cells were infected with equal amounts of virus, DNA was isolated 5 hours post infection (hpi) and RT-qPCR was performed to measure MLV DNA levels. As expected, both F-MLV WT and F-MLV/Ampho^Env^ had similar MLV DNA levels in the infected XC cells independent of the presence of mSERINC5 (Fig. 9A, B). In the case of gGag^mut^F-MLV, virus DNA levels were similar in XC cells regardless of the presence of mSERINC5 in the virions (Fig. 9A), similar to what we observed when we infected Mus Dunni cells (Fig. 3E). However, when infecting XC cells with gGag^mut^F-MLV/Ampho^Env^, we observed that viral DNA levels were reduced when the virus was produced in the presence of mSERINC5 (Fig. 9C). These results are similar to what we observed before when we used Mus Dunni cells (Fig. 5F). Thus, the above findings suggest that the resistance of ecotropic MLV to mSERINC5 is independent of the route of virus entry as previously hypothesized.

**Fig. 9.**
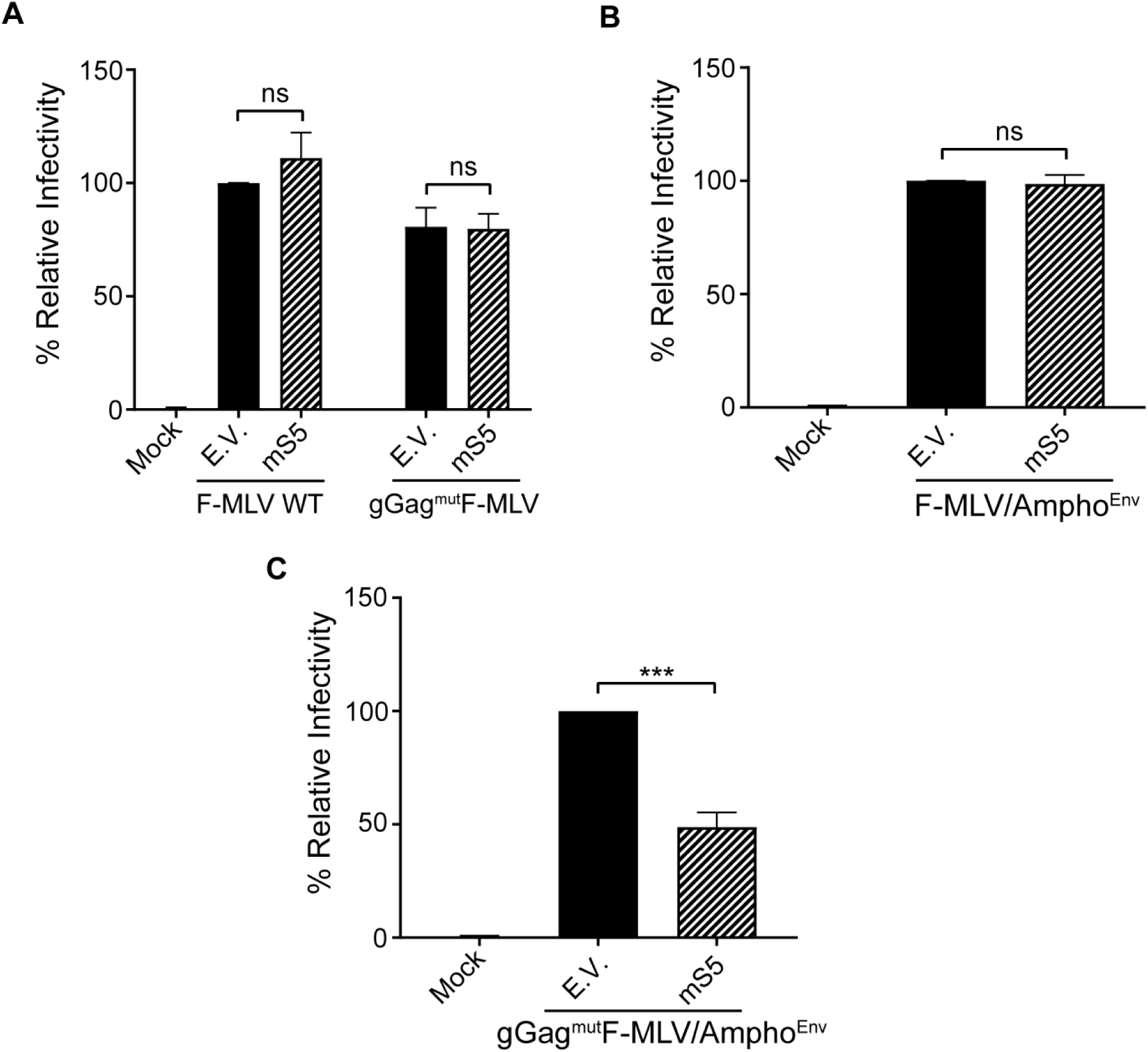
MLV sensitive to mSERINC5 restriction is independent of the route of virus entry. (A-C) XC cells were infected with equal amounts of F-MLV WT (A), gGag^mut^F-MLV (A), F-MLV/Ampho^Env^ (B), and gGag^mut^F-MLV/Ampho^Env^ (C) produced in the presence of mSERINC5 or empty vector. Cells were harvested 5 hours post infection and MLV DNA levels were measured by RT-qPCR and normalized to GAPDH. Percentage (%) of relative infectivity was determined with respect to virus generated in the presence of empty vector. All results are presented as mean ± SD. Statistical significance was determined by unpaired t test (two-tailed). ***, p < 0.001; ns, not significant. Results are shown for n = 3 independent experiments. (mouse SERINC5, mS5; empty vector, E.V.)

## DISCUSSION

*In vitro* studies have shown that SERINC5 can block retrovirus infection (3-5, 25). However, many retroviruses have developed mechanisms that can counteract SERINC5-mediated restriction. In the case of MLV, glyco-Gag inhibits SERINC5 restriction by removing SERINC5 from the plasma membrane and thus blocking its incorporation in the budding virions (25, 31). While a lot of work has been done to understand the *in vitro* antiretroviral functions of SERINC proteins (3-5, 25, 43), nothing is known about the antiretroviral function of this family of proteins *in vivo*. For many antiretroviral proteins, e.g. SAMHD1, their antiviral effect *in vitro* is not always recapitulated *in vivo* (27). Many times, *in vitro* experiments are done using cell types that are not naturally infected by retroviruses, or SERINC proteins are expressed significantly higher than physiological levels. Finally, natural MLV infection in mice involves a multitude of different cell types, something that cannot be easily examined *in vitro*.

In order to examine whether SERINC3 and SERINC5 have antiretroviral properties *in vivo*, we generated SERINC5-/- mice and confirmed the disruption of the gene by RT-qPCR, as there is no antibody available to detect endogenous mSERINC5 protein levels. SERINC5-/- mice had no apparent defects and, thus, are a useful model to examine the *in vivo* effect of mSERINC5 on retrovirus infection. Using wild type and glyco-Gag mutant F-MLV, we saw no differences in viral titers in the spleens of either the SERINC5-/- or the BL/6 infected mice. On the other hand, we observed glyco-Gag dependent exclusion of mSERINC3 and mSERINC5 from the budding virions. We speculated that the lack of any obvious antiviral effect may be because the viruses used in these experiments had the ecotropic envelope, which makes them resistant to SERINC5 restriction (5, 43). Previous reports, using pseudoviruses, showed that the presence of amphotropic or xenotropic MLV envelope on the surface of virions rendered them susceptible to human SERINC5 restriction *in vitro* (5, 33). To examine the effect of the envelope on mSERINC5 restriction in fully infectious virus clones, we generated chimeric F-MLV constructs, where the ecotropic envelope was replaced with that of amphotropic MLV. Using these chimeric viruses, we found that the glyco-Gag mutant MLV with an amphotropic envelope replicated at higher levels in the spleens of SERINC5-/- mice compared to BL/6 mice, demonstrating the critical role of the viral envelope on mSERINC5 restriction *in vivo*. This report provides the first direct evidence that mSERINC5 is an antiretroviral factor *in vivo* and that the retroviral envelope is important in determining *in vivo* sensitivity to mSERINC5 restriction. Of note is the striking difference in the glyco-Gag requirement for virus infectivity between viral particles with the amphotropic envelope and those with ecotropic envelope. While the former required glyco-Gag for infectivity both *in vivo* and *in vitro*, the latter were unaffected *in vitro*, while *in vivo* there was only about a 10-fold drop in infectivity when glyco-Gag was mutated. It is not clear why the presence of different envelopes modulates the role of glyco-Gag on virus infectivity. The relationship between envelope and glyco-Gag needs to be elucidated. To circumvent the absence of an antibody that detects endogenous mSERINC5, we compared the infectivity between viral particles with the amphotropic envelope and mutated glyco-Gag isolated from the spleens of either SERINC5-/- or BL/6 infected mice. We found that virions isolated from the SERINC5-/- mice were more infectious than those derived from BL/6 mice, which suggests that mSERINC5 is incorporated in virions *in vivo* and blocks virus infectivity. This is the first evidence that suggest mSERINC5 is incorporated in viral particles *in vivo* and can affect virus infectivity. Furthermore, it has been hypothesized in the field that the route of entry of a virus could affect SERINC5 restriction. Our data suggest that route of entry of a virus does not affect susceptibility to SERINC5 restriction, as ecotropic MLV was resistant to SERINC5 restriction in both Mus Dunni cells, where the virus enters in a pH-dependent manner, and in XC cells, where entry occurs independent of pH. Thus, another factor and not the route of entry of a virus affects the susceptibility of the retrovirus envelope to SERINC5.

Similar to mouse APOBEC3 (24, 38, 41, 50), our findings show that mSERINC5 potently restricts MLV infection *in vivo*. SERINC5 and mouse APOBEC3 restrict retrovirus infection at different steps of the retroviral life cycle, SERINC5 during virus-cell fusion (7), while APOBEC3 during reverse transcription (42, 50). Using SERINC5/APOBEC3 double knockout mice we show that they indeed act synergistically to block retrovirus infection. This is the first time that two different antiretroviral factors have been shown to act in unison *in vivo*. However, how these two factors come together to synergistically restrict MLV infection *in vivo* needs further investigation.

We also examined the role of mSERINC3 on retrovirus infectivity *in vivo* and *in vitro*. Human SERINC3 has been shown to be antiretroviral, albeit not as potent as SERINC5 (3, 4), while there are differing opinions on the antiretroviral role of mSERINC3 (25, 51). Our *in vitro* and *in vivo* findings show that mSERINC3 has no antiretroviral functions against MLV, although it is targeted for internalization by glyco-Gag *in vitro*. It is possible that the antiretroviral function of the human homologue of SERINC3 was acquired at a later point in the evolutionary scale possibly due to selective pressure from a retrovirus that was absent in the mouse evolutionary history or it is possible that the murine SERINC3 has different viral targets than the human homologue.

## MATERIALS AND METHODS

### Ethics Statement

All mice were housed and bred according to the policies of the Institutional Animal Care and Use committee of the University at Buffalo. The experiments performed with mice in this study were approved by this committee (MIC 46038Y).

### Generation of SERINC5-/- mice

To generate the SERINC5-/- mice, exon1 and exon 2 were targeted by 2 guide RNAs (gRNAs) (gRNA1:5’- ATGTCTGCCCGGTGCTGTGCTGG and gRNA2: 5’ GCCCTAAGTTCCGACAGTCTCGG) using the IDT Alt-R crRNA/trRNA CRISPR/Cas9 technology. The gRNAs and recombinant Cas9 (RNPs) were electroporated into zygotes of C57BL/6N mice (Charles River Laboratories). DNA from the mice generated from the electroporations was sent to the Genome Engineering and iPSc Center at Washington University in St. Louis were they were screened via targeted NGS for the presence of deletion or indels at the target site. One founder line was discovered to have a sizable 49kb deletion between exon 1 and 2. As such, for the purpose of genotyping we used two sets of primers SEREx1F and SEREx2R / SEREx1F and SERInt2R (see Table S1); the first set of primers amplifies the deleted allele as the two primers are 49kB apart in the BL/6 mice, the second set of primers amplifies a region of about 250bp.

### Generation of SERINC5/APOBEC3 -/- mice

APOBEC3 mice were bred at SUNY, University at Buffalo as previously described (11, 52). The SERINC5-/- mice described in this study were crossed with the APOBEC3-/- mice to generate SERINC5/APOBEC3-/- mice.

### Generation of SERINC3-/- mice

SERINC3-/- mice were derived from *in vitro* fertilization of C57BL/6N females with sperm from a single heterozygous male from the mutant mouse line C57BL/6NTac-SERINC3^tm1a(KOMP)Wtsi^/WtsiFlmg constructed by the Wellcome Trust Sanger Institute Mouse Genetics Project (Sanger MGP). Cre-Deleter mice (C57BL/6NTac-Gt(ROSA)26Sor^tm16(cre)Arte^), where the cre gene is expressed under the control of the Gt(ROSA)26Sor gene, were purchased from Taconic Biosciences and used to develop SERINC3-/- mice. Deletion by cre recombinase in the SERINC3^tm1a(KOMP)Wtsi^/WtsiFlmg results in the removal of exon3 leading to a frameshift mutation and a premature stop codon. The following primers were used for genotyping: Serinc3_35556_F and Serinc3_35556_R / SERINC3Ex3F and SERINC3Ex3R for detection of WT allele, Serinc3_35556_F and CAS_R1_Term / DelF and DelR for detection of deleted allele. Primers used for detection of cre were CreF and CreR (see Table S1).

### Cell culture and transfection

293T (ATCC) and 293FT cells (Invitrogen) were cultured in Dulbecco’s Modified Eagle Media (DMEM; Gibco) with 10% (vol/vol) fetal bovine serum (FBS; Sigma), 0.1 mM non-essential amino acids (Gibco), 6 mM L-glutamine (Gibco), 100 mg/ml penicillin and streptomycin (P/S; Gibco), 1 mM sodium pyruvate (Gibco), and 500 μg/ml geneticin (Gibco). NIH3T3 cells and XC cells (ATCC) were cultured in DMEM with 10% FBS and P/S. Mus Dunni cells were maintained in RPMI media with 10% FBS and P/S. EL4 cells were cultured in RPMI with 10% FBS, P/S, and 0.05 mM β-mercaptoethanol (βME; Bio-Rad). MutuDC1940 cells (30) were cultured in IMDM with 8% FBS, 100 mg/ml P/S, 1 mM sodium pyruvate, 10 mM HEPES (Corning), and 0.05 mM βME. All transfections were performed using the Lipofectamine 3000 transfection kit (Invitrogen) per manufacturer’s recommendation.

### Plasmids

The F-MLV WT provirus construct pLRB302 has been previously described (53, 54). To generate glyco-Gag-deficient F-MLV (gGag^-^F-MLV) provirus, two stop codons were introduced at residues 32 and 55 of the glyco-Gag reading frame using the Phusion SDM kit (ThermoFisher Scientific). The primers used to introduce the changes (underlined) were: 5’-TGCACCCCCCT<u>A</u>AGAGGAGGGGT-3’/ 5’-CCAAAGAGTCCAAAACGATCGGGATGG-3’ and 5’-GTCTGAGTTTT<u>A</u>GCTTTCGGTTTGGAA-3’/ 5’-GGGGGCGGAAACCGTTTTAGCC-3’. To minimize polymerase-driven errors, XhoI-EcoRI fragment from gGag^-^F-MLV construct was subcloned into F-MLV WT backbone. Glyco-Gag mutated F-MLV (gGag^mut^F-MLV) was generated by mutating P31, Y36, L39 and R63 residues of glyco-Gag to alanine by site directed mutagenesis using the Phusion SDM kit (ThermoFisher Scientific) (primer pairs used: 5’-CTCTTTGGTGCACCC<u>G</u>CCTTAGAGGAGGGG<u>GC</u>TGTGGTT<u>GC</u>GGTAGGAGACAGAGG-3’/ 5’-TCCAAAACGATCGGGATGGTTGGACTC-3’ for P31A, Y36A, and L39A and 5’-CTTTCGGTTTGGAACCGAAGC<u>GC</u>CGCCGCGCGTCTTGTCTGC-3’/ 5’-CAAAAACTCAGACGGGGGCGGAAAC-3’ for R63A). A plasmid containing the amphotropic MLV *env* gene construct (pSV-A MLV env) was obtained from NIH AIDS Reagent program, Division of AIDS, NIAID, NIH: SV-A-MLV-env from Dr. Nathaniel Landau and Dr. Dan Littman (55). F-MLV amphotropic envelope chimeras (F-MLV/Ampho^Env^, gGag^-^F-MLV/Ampho^Env^, and gGag^mut^F-MLV/Ampho^Env^) were created by replacing the entire *env*-coding sequence of F-MLV WT with amphotropic *env*-coding sequence from p-SV-A MLV *env* using NEBuilder HiFi DNA Assembly Kit (New England Biolabs). The primers used for swapping the *env*-coding regions were: 5’-ATAAAAGATTTTATTTAGTTTCCAGAAAAAGGGGGGA-3’/ 5’-CGTTGAACGCGCCATGTCGATTCCGATGGTGGCTC-3’ for the F-MLV backbone and 5’-ACCATCGGAATCGACATGGCGCGTTCAACGCTCTC-3’/ 5’-AATAAAATCTTTTATCTATGGCTCGTACTCTATAGG-3’ for the amphotropic MLV *env*. pBJ5-hSERINC3-HA and pBJ5-hSERINC5-HA were obtained from Heinrich Gottlinger(4). pBJ5-mSERINC3-HA and pBJ5-mSERINC3-HA were generated by replacing the human SERINC genes with the murine ones. mSERINC3/mSERINC5 were PCR amplified from mSERINC3 and mSERINC5 cDNAs (Dharmacon Incorportated) using the primers 5’-GGGCTCGAGATGGGGGCCGTCCTCG-3’/ 5’-CCCGCGGCCGCTTAAGCGTAATCTGGAACATCGTATGGGTAGCTGAAGTCCCGACCT GTG-3’ for mSERINC3 and the primers 5’-GGGCTCGAGATGTCTGCCCGGTGCTGTG-3’/ 5’-CCCGCGGCCGCTTAAGCGTAATCTGGAACATCGTATGGGTACACAGAGAACTGCCT GGA-3’ for mSERINC5 followed by XhoI/NotI digestion and ligation in the pBJ5 vector. All constructs generated were confirmed by DNA sequencing. Primers used for sequencing are listed in Table S2.

### Virus preparation for animal experiments

Virus stocks were prepared by transfecting 293FT cells (Invitrogen) seeded in 10 cm cell culture dishes with the viral constructs described above. Culture supernatants were harvested 48 h after transfection, filtered and treated with 10 U/ml DNase I (Roche) for 40 min at 37 °C. Titers of viruses were determined by infecting Mus Dunni cells for 48 h. For the ecotropic MLV viruses, immunofluorescent foci assays (IFAs) were performed to count infectious centers (ICs) using Mab720 (56) followed by DAB staining with horseradish peroxidase (HRP) substrate (Vector laboratories). In the case of the chimeric viruses with the amphotropic MLV envelope, IFAs were performed using the Mab573 (57) followed by immunostaining with Alexa Fluor™ 488-conjugated goat anti-mouse antibody (Invitrogen); samples were then processed using the BioTek Cytation 1 Imaging Plate Reader (BioTek).

### Detection of SERINC3/5 incorporation in virions by immunoblotting

0.5 × 10^6^ 293T cells (Invitrogen) were seeded in 6-well culture plates and co-transfected with viral DNA construct (5 μg) and either pBJ5-mSERINC3-HA or pBJ5-mSERINC5-HA or empty vector (100 ng). Cells and culture media were harvested 48 h after transfection. Cells lysates were prepared as previously described (6). Briefly, cells were lysed in DM lysis buffer (0.5% (w/v) n-Decyl-β-D-maltopyranoside in 20 mM Tris-HCl, pH 7.5, 10 % (v/v) glycerol, 1× protease inhibitors and 25 U/ml Benzonase), centrifuged at 14,000 rpm followed by the collection of the supernatant fraction. Culture media were pelleted through a 30% sucrose cushion as previously described (11). Cell lysates or virus pellets were mixed with 1× sample loading buffer and incubated at 37 °C for 15 min before resolving them on 10% sodium dodecyl sulfate polyacrylamide gel electrophoresis (SDS-PAGE) gels. Blots were probed using the following antibodies: Polyclonal goat anti-MLV (11) (NCI repository), rat anti-MLV p30 R187 (ATCC® CRL-1912™), rabbit anti-HA (Cell Signaling Technology), monoclonal anti-β-actin (Sigma-Aldrich), HRP-conjugated anti-rabbit (Cell Signaling Technology), -anti-rat (Cell Signaling Technology), -anti-mouse (EMD Millipore) and -anti-goat (Sigma Aldrich) were used for detection using the enhanced chemiluminescence detection kits Clarity and Clarity Max ECL (Bio-Rad).

### Target cell assays

Virions were prepared and treated with 10 U/ml DNase I (Roche) for 40 min at 37 °C as described above. Virus preparations were normalized following quantitation by MLV-p30 immunoblotting and analysis using ImageJ software (National Institutes of Health; https://imagej.nih.gov/ij/). Equal amounts of viruses were used to infect 0.8 × 10^5^ Mus Dunni or XC sarcoma cells in a 6-well plate. Cells were harvested 5 hpi. DNA was extracted using the DNeasy Blood and Tissue kit (Qiagen) per manufacturer’s instructions and RT-qPCR was performed using the Power SYBR Green PCR master mix kit (Applied Biosystems). The MLV primers used were for the ecotropic MLV infections (5’-TACAGGGAGCTTACCAGGCA-3’/ 5’-GTTCCTATGCAGAGTCCCCG-3’) and for the amphotropic envelope chimeric MLV infections (5’-AAGTCCAAGTGTCCCACAGC-3’/5’-AGCCCACATTGTTCCGGC-3’). GAPDH primers (5’-CCCCTTCATTGACCTCAACTACA-3’/ 5’-CGCTCCTGGAGGATGGTGAT-3’) were used for normalization. The CFX384 Touch Real-Time PCR detection system (Bio-Rad) was used for all RT-qPCR assays described in this study. The reactions were performed under the following conditions: 95 °C for 10 min, followed by 40 cycles of 95 °C for 10 s and 60 °C for 30 s. The relative amplification for each sample was quantified from standard curves generated using known quantities of DNA standard templates.

### *In vivo* infections

For all SERINC5 and SERINC3 *in vivo* experiments, newborn mice were infected by intraperitoneal (IP) injections with 5 × 10^3^ ICs of virus as previously described (11, 24, 38). Spleens from infected mice were harvested 10 days post-infection and used for infectivity assays described below.

### Infectivity assays

MLV infection levels in the spleens were measured by IC assays as previously described (38). Briefly, Mus Dunni cells were seeded at a density of 0.8 × 10^5^ per well of a 6-well plate. Next day, the cells were co-cultured with 10-fold dilutions of splenocytes isolated from infected mice. ICs were determined 48 hpi after performing IFAs, as described above (**Virus preparation for animal experiments**).

### Growth curve experiment

NIH3T3 cells (40,000/well) were seeded in a 6-well plate. Cells were infected the next day with the different viruses used in this project (0.1 MOI). After 2 h infection, cells were washed 3 times with PBS and cultured in 2 ml media for 5 days. Culture supernatants and cells were harvested each day and stored at –80 °C. Concentrations of MLV p30 in the supernatants were determined using QuickTiter™ MuLV Core Antigen ELISA Kit (MuLV p30; Cell Biolabs). DNA was isolated from the infected cells using the DNeasy Blood and Tissue Kit (Qiagen) and viral DNA levels were determined by RT-qPCR, as described above (**Target cell assays**).

### SERINC3/5 transcription levels

Tissues were harvested from 3 month old BL/6, SERINC5-/- and SERINC3-/- mice and total RNA was isolated using Trizol (Invitrogen) as per manufacturer’s instruction. cDNA was synthesized using the SuperScript III First Strand Synthesis kit (Invitrogen). RT-qPCR was performed as described above (**Target cells assays**). The primers used: mSERINC3 5’-TGACCAATGAACCTGAGCGG-3’/ 5’-GCTGTTGCTCGAAGTACGGA-3’, and mSERINC5 5’-GTCCAGAATCGACAGCCACA-3’/ 5-ATCCGAGCTCGACCTTGTTG-3’. Actin primers (5’-TGGAATCCTGTGGCATCCATGAAAC-3’/ 5’-TAAAACGCAGCTCAGTAACAGTCCG -3’) were used for normalization.

MutuDC1940 and EL4 cells were infected with MLV by the spinoculation method as previously described (38). Briefly, cells were seeded in a 96-well plate at a cell density of 0.8 × 10^5^/100 μl of culture media. Next day, the cells were infected with MLV (0.1 MOI) and the plate was centrifuged at 1,800 rpm for 2 h at room temperature. After centrifugation, the cells were washed 3 times with PBS and maintained in 100 μl of media at 37°C. Cells were harvested at different time points. RNA was isolated and SERINC3/5 expression levels were determined by RT-qPCR as described above (**Target cell assays**).

### FACS analysis and sorting

Blood was collected via retro-orbital bleeding from SERINC5-/-, SERINC3-/- and BL/6 mice of similar ages. PBMCs were stained with anti-CD3-APC (BD Biosciences), anti-CD45R/B220-FITC (BD Biosciences), anti-CD11c-APC (BD Biosciences) and anti-mouse F4/80-FITC (Biolegend) (23). Stained cells were processed using BD LSRFortessa. Results were analyzed using FlowJo software.

B cells, T cells, and DCs were sorted from blood using a BD FACSAria Fusion Cell Sorter. SERINC1/2/3/4/5 expression profile was determined by RT-qPCR as described above (**Target cells assays**). In addition to the SERINC3/5 and GAPDH primers mentioned above (**SERINC3/5 transcription levels**), the following primers were used to detect the other SERINC family members: SERINC1 (5’-CTGTTCAGTGGTGGCATCCTCA-3’/ 5’-CCTGACTGTTGTTGGAGGTACG-3’), SERINC2 (5’-AATCAGCGGTGGCTGTGTAAGG-3’/ 5’-ACATCAGTGCCACAGCAGCGAT-3’) and SERINC4 (5’-CCGCAAGCAACCAAACTCTG-3’/ 5’-CTGCTACTGAGGTATCCGGTATT-3’).

### Infectivity measurement of *in vivo-*derived virus

Two day old SERINC5 -/- and BL/6 mice were infected by IP injections with 4 × 10^5^ ICs of virus and spleens were harvested 16 days post-infection. Spleen extracts were prepared as previously described (58) with some modifications. Briefly, spleens were mashed through a cell strainer and splenocytes were collected in 5 ml (for F-MLV/Ampho^Env^ virus) or 2 ml (for gGag^mut^F-MLV/Ampho^Env^ virus) of RPMI with 10% FBS and 100 mg/ml P/S. Cells were pelleted at 1,200 rpm for 10 minutes at 4 °C. The supernatant was removed and centrifuged at 3,000 rpm for 10 minutes at 4 °C and then filtered through a 0.45 μm syringe filter. RNA was isolated from 100 μl of the supernatant using the RNeasy kit (Qiagen) per manufacturer’s recommendation. cDNA synthesis and RT-qPCR were performed as described above (**SERINC3/5 transcription levels**). Normalized spleen extracts (based on RNA copies) were used to infect 0.3 × 10^5^ Mus Dunni cells seeded in a 24-well plate. Cells were harvested 19 hpi and the amount of viral DNA in the infected cells was determined by RT-qPCR, as described above (**Target cell assays**).

### Ammonium chloride treatment

One day prior to infection, NIH3T3, Mus Dunni or XC cells were seeded in a six well plate (0.8 × 10^5^ cells/ well). Cells were pretreated with 30 mM ammonium chloride (NH_4_Cl) for 30 minutes before infection with F-MLV WT or F-MLV/Ampho^Env^ viruses (MOI 0.5) (45). The same concentration of NH_4_Cl was maintained during the infection period and an additional 2 h, after which cells were maintained in fresh media without NH_4_Cl. Cells were harvested 19 hpi and the amount of viral DNA in the infected cells was determined by RT-qPCR, as described above (**Target cell assays**).

### Statistical analysis

Statistical analyses were performed using GraphPad Prism software version 8.2. Statistical tests used to determine significance are described in the figure legends. A difference was considered to be significant if p-value was < 0.05.

## ACKNOWLEDGEMENTS

We thank Susan Ross, Leonard “Pug” Evans and Heinrich Gottlinger for kindly providing us with reagents that were used in this study. We also thank Ray Kelleher for help with the primary cell sorting and the Gene Targeting and Transgenic Shared Resource at Roswell Park Comprehensive Cancer Center for assisting in the derivation of the SERINC5-/- and SERINC3-/- mice. We thank the Welcome Trust Sanger Institute Mouse Genetics Project (Sanger MGP) and its funders for providing the mutant mouse line (C57BL/6NTac-SERINC3^tm1a(KOMP)Wtsi^/WtsiFlmg) and INFRAFRONTIER/EMMA (www.infrafrontier.eu). Funding information may be found at www.sanger.ac.uk/mouseportal and associated primary phenotypic information at www.mousephenotype.org. Finally, we would like to thank Sabine Baxter at the Cell Center Services of the University of Pennsylvania for assisting us with our antibody purification needs.

The work was supported by National Institutes of Health grant R21-AI144147-01A1 and a Mathilde Krim Fellowship in Basic Biomedical Research Phase II, American Foundation for AIDS Research 109741-63-RKHF.

## Notes

### Competing Interest Statement

The authors have declared no competing interest.

