## Supplemental Data for "SERINC5 potently restricts retrovirus infection *in vivo*"

**This file includes:**

Tables S1 to S2

Figures S1 to S2

### Supplemental Data

**Table S1. Primers used for genotyping the mice**

| S. No. | Name of primer | Nucleotide sequence |
|---|---|---|
| 1. | SEREx1F | 5’- GGGGATAGGAAGCAAGACGA-3’ |
| 2. | SEREx2R | 5’- TGCTTCATCACAGAGGGTGTC-3’ |
| 3. | SERInt2R | 5’-TCCTGGATTTATCCCACAGG-3’ |
| 4. | Serinc3_35556_F | 5’-CCCCACTGTCTGAACAAACG-3’ |
| 5. | Serinc3_35556_R | 5’-TTCCTAACACGCACATGGTTG-3’ |
| 6. | SERINC3Ex3F | 5’-CAGAGAAAGATTGTGACGTGCTG-3’ |
| 7. | SERINC3Ex3R | 5’-TGCTGCTCTGGGATCTTTACTTG-3’ |
| 8. | CAS_R1_Term | 5’-TCGTGGTATCGTTATGCGCC-3’ |
| 9. | CreF | 5’-GAACCTGATGGACATGTTCAGG-3’ |
| 10. | CreR | 5’-AGTGCGTTCGAACGCTAGAGCCTGT-3’ |
| 11. | DelF | 5’-TCCCCCTGAACCTGAAACATAA-3’ |
| 12. | DelR | 5’-TGATTTGAACTGATGGCGAGC-3’ |

**Table S2. Primers used for sequencing of proviral constructs**

| S. No. | Name of primer | Nucleotide sequence |
|---|---|---|
| 1. | FMLV gGag Sequencing_F1 | 5’-TTCGGGGGCCATTTTTGTGG-3’ |
| 2. | FMLV gGag Sequencing_R | 5’-AGCCTGGCCCATGTTTTCAG-3’ |
| 3. | FMLV gGag Sequencing_F2 | 5’-CTGGCGGATCCGTGGTGGAAC-3’ |
| 4. | FMLV Env Sequencing_F | 5’-TCTCAAAGTGGACGGCATTG-3’ |
| 5. | FMLV Env Sequencing_R | 5’-TTACTGCGGCTATCAGGCTAAGC-3’ |

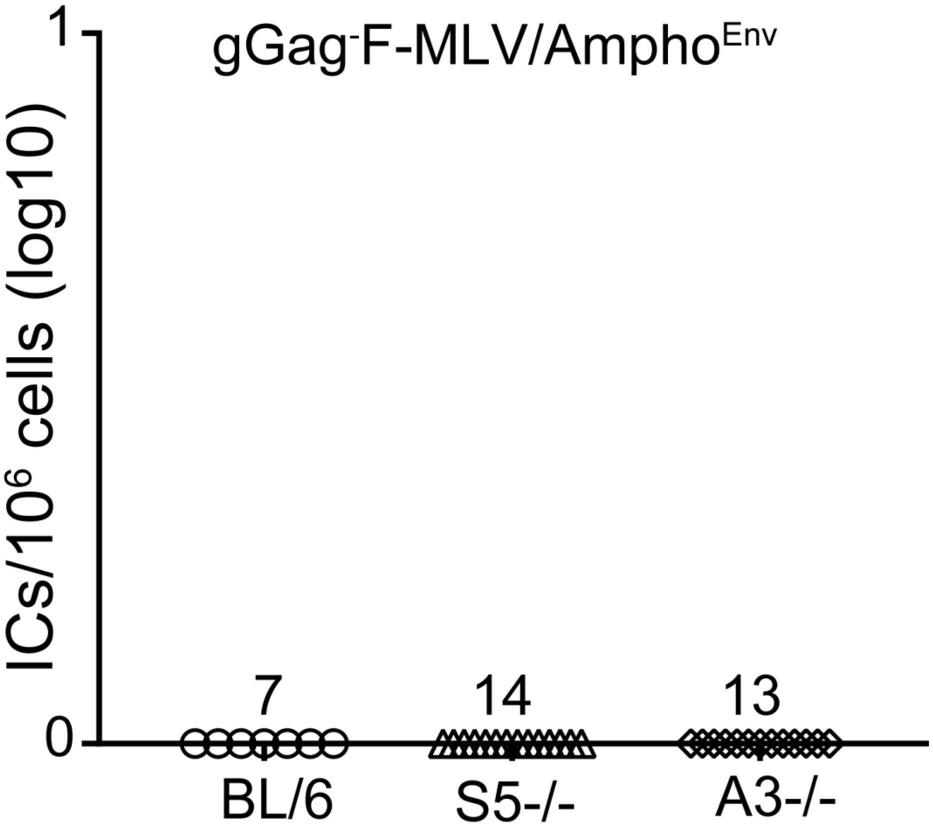


Fig. S1 Glyco-Gag deleted Amphotropic MLV does not replicate *in vivo*. Newborn mice were infected with $gGag^{-}$F-MLV/$Ampho^{Env}$ and virus titers in spleens were measured 10 days post infection. Each point represents the titer obtained from an individual mouse and the mean for each group is shown by a horizontal line. Mice were derived from 3-4 litters each; the C57BL/6 and SERINC5-/- are the littermates. Numbers of mice used in each group are indicated in the X-axis. (SERINC5, S5; C57BL/6, BL/6; APOBEC3-/-, A3-/-; Infectious centers, ICs)

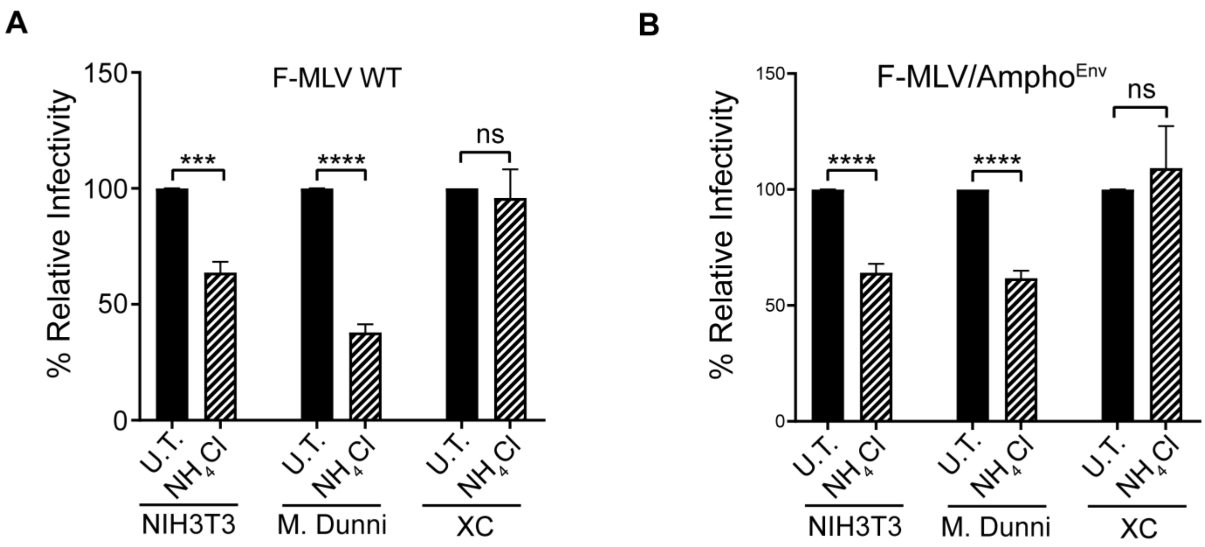


Fig. S2 pH dependency of MLV infection is cell-type dependent. (A, B) NIH3T3, Mus Dunni and XC cells were infected with F-MLV WT (A) and F-MLV/Ampho$^{Env}$ (B) (MOI 0.5) in the presence or absence of 30 mM ammonium chloride. Cells were harvested 19 hours post infection and MLV DNA levels were measured by RT-qPCR and normalized to GAPDH. Percentage (%) of relative infectivity was determined with respect to untreated control (U.T.). All results are presented as mean ± SD. Statistical significance was determined by unpaired t test (two-tailed). ***, $p < 0.001$; ****, $p < 0.0001$; ns, not significant. Results are shown for n = 3 independent experiments. (Ammonium chloride, $NH_4Cl$; untreated, U.T.)
